# Rare earth element (REE)-dependent growth of *Pseudomonas putida* KT2440 depends on the ABC-transporter PedA1A2BC and is influenced by iron availability

**DOI:** 10.1101/670216

**Authors:** Matthias Wehrmann, Charlotte Berthelot, Patrick Billard, Janosch Klebensberger

## Abstract

In the soil-dwelling organism *Pseudomonas putida* KT2440, the rare earth element (REE)-utilizing and pyrroloquinoline quinone (PQQ)-dependent ethanol dehydrogenase PedH is part of a periplasmic oxidation system that is vital for growth on various alcoholic volatiles. Expression of PedH and its Ca^2+^-dependent counterpart PedE is inversely regulated in response to lanthanide (Ln^3+^) bioavailability, a mechanism termed the REE-switch. In the present study, we demonstrate that copper, zinc, and in particular, iron availability influences this regulation in a pyoverdine-independent manner by increasing the minimal Ln^3+^ concentration required for the REE-switch to occur by several orders of magnitude. A combined genetic- and physiological approach reveals that an ABC-type transporter system encoded by the gene cluster *pedA1A2BC* is essential for efficient growth with low (nanomolar) Ln^3+^ concentrations. In the absence of *pedA1A2BC*, a ~100-fold higher La^3+^ concentration is needed for PedH-dependent growth but not for the ability to repress growth based on PedE activity. From these results, we conclude that cytoplasmic uptake of lanthanides through PedA1A2BC is essential to facilitate REE-dependent growth under environmental conditions with poor REE bioavailability. Our data further suggest that the La^3+^/Fe^3+^ ratio impacts the REE-switch through the mismetallation of putative La^3+^-binding proteins, such as the sensor histidine kinase PedS2, in the presence of high iron concentrations. As such, this study provides an example for the complexity of bacteria-metal interactions and highlights the importance of medium compositions when studying physiological traits *in vitro* in particular in regards to REE-dependent phenomena.

## Introduction

Metal ions are essential for all living organisms as they play important roles in stabilizing macromolecular cellular structures, by catalyzing biochemical reactions or acting as cofactors for enzymes (Gray, 2003; Merchant and Helmann, 2012). They can, however, also be toxic to cells at elevated levels through the generation of reactive oxygen species or by aspecific interactions such as mismetallation (Cornelis et al., 2011; Dixon and Stockwell, 2014; Foster et al., 2014). Bacteria have hence developed a sophisticated toolset to maintain cellular metal homeostasis (Andrews et al., 2003; Chandrangsu et al., 2017; Schalk and Cunrath, 2016; Semrau et al., 2018). Common mechanisms include release of metal-specific scavenger molecules, the activation of high-affinity transport systems, the production of metal storage proteins, and the expression of specific efflux pumps.

As it is the case for all strictly aerobic bacteria, the soil-dwelling organism *Pseudomonas putida* KT2440 has a high demand for iron but faces the challenge that the bioavailability of this metal is very poor under most toxic environmental conditions due to the fast oxidation of Fe^2+^- and the low solubility of Fe^3+^-species (Andrews et al., 2003). One strategy of many bacteria to overcome this challenge is to excrete self-made peptide-based siderophores (such as pyoverdines) into the environment that bind Fe^3+^ with high affinity, and thereby increase its bioavailability (Baune et al., 2017; Cornelis and Andrews, 2010; Salah El Din et al., 1997). A second adaptation of *P. putida* cells to iron-limitation is a change in the proteomic inventory in order to limit the use of Fe-containing enzymes, exemplified by the switch from the Fe-dependent superoxide dismutase (SOD) to a Mn-dependent isoenzyme or by re-routing of entire metabolic pathways (Kim et al., 1999; Sasnow et al., 2016). In contrast, when Fe bioavailability is high, the production of the bacterioferritins Bfrα and Bfrβ is increased to enable intracellular storage and thereby improve cellular fitness under potential future conditions of iron starvation (Chen et al., 2010). The regulatory mechanisms for metal homeostasis of *P. putida* cells in response to other essential metal ions such as Co, Cu, Mg, Mo, Ni, and Zn are less well explored. Genes encoding for transport systems associated with the uptake and efflux of these metals can, however, be found in its genome (Belda et al., 2016; Nelson et al., 2002), and some of these have been studied in more detail (Miller et al., 2009; Ray et al., 2013).

We have recently reported that *P. putida* KT2440 is capable of using rare earth elements (REEs) of the lanthanide series (Ln^3+^) when growing on several alcoholic substrates (Wehrmann et al., 2017, 2019). Under these conditions, the cells use the pyrroloquinoline quinone (PQQ)-dependent ethanol dehydrogenase (EDH) PedH, to catalyze their initial oxidation within the periplasm. Like many other organism, *P. putida* harbors an additional, Ln^3+^-independent functional homologue of PedH termed PedE that depends on a Ca^2+^ ion as metal cofactor (Takeda et al., 2013; Wehrmann et al., 2017). Depending on the availability of REEs in the environment, *P. putida* tightly regulates PedE and PedH production (Wehrmann et al. 2017, 2018). In the absence of Ln^3+^, growth is solely dependent on PedE whereas PedH transcription is repressed. The situation immediately changes in the presence of small amounts of Ln^3+^ (low nM range) leading to a strong induction of the Ln^3+^-dependent enzyme PedH and repression of its Ca^2+^-dependent counterpart PedE. For *P. putida* KT2440 the PedS2/PedR2 two component system (TCS) is a central component of this inverse regulation (Wehrmann et al., 2018). Notably, the REE-switch in *P. putida* was also found to be influenced by environmental conditions, as the critical La^3+^ concentrations required to support PedH-dependent growth differs dramatically depending on the medium used, ranging from 5 nM up to 10 μM (Wehrmann et al., 2017). Lanthanides are only poorly available in natural environments (often picomolar concentrations) due to the formation of low soluble hydroxide and/or phosphate (Firsching and Brune, 1991; Meloche and Vrátný, 1959). The presence of active uptake systems to facilitate REE-dependent growth in bacteria has thus been favored by many researchers (Aide and Aide, 2012; Cotruvo et al., 2018; Gu et al., 2016; Gu and Semrau, 2017; Markert, 1987; Picone and Op den Camp, 2019; Tyler, 2004). A transcriptomic study of *M. trichosporium* OB3b cells observed that multiple genes encoding for different active transport systems were among the most regulated in the presence of cerium (Gu and Semrau, 2017). In addition to these findings, it has been found that a specific TonB-dependent receptor protein as well as a TonB-like transporter protein are highly conserved in bacteria that carry genes encoding for Ln^3+^-dependent MDHs (Keltjens et al., 2014; Wu et al., 2015). Only very recently, different studies indeed identified both an ABC-transporter and TonB-dependent receptor proteins that are needed for REE-dependent growth of methano- and methylotrophs, strongly suggesting the existence of an uptake system that specifically transports a Ln^3+^-chelator complex in these organisms (Groom et al., 2019; Ochsner et al., 2019; Roszczenko-Jasińska et al., 2019).

With the present study, we show that a homologous ABC-transporter system, encoded by the gene cluster *pedA1A2BC*, is also essential for lanthanide-dependent growth in the non-methylotrophic organism *Pseudomonas putida* KT2440 under low (nanomolar) concentrations of REEs. Notably, no homolog of the TonB-dependent receptor proteins found in methanotrophic or methylotrophic strains could be identified within the genome of *P. putida* KT2440 indicating either a lack or substantial differences in the chemical nature of such a Ln^3+^-specific chelator system. Finally, we show that the siderophore pyoverdine plays no essential role for growth under low REE concentrations but provide compelling evidence that in addition to Cu^2+^ and Zn^2+^ the Fe^2+/3+^ to Ln^3+^ ratio can significantly alter the REE-switch most likely through mismetallation.

## Materials and methods

### Bacterial strains, plasmids and culture conditions

The *E. coli* and *P. putida* KT2440 strains and the plasmids used in this study are described in **Table 1**. Maintenance of strains was routinely performed on solidified LB medium (Maniatis et al., 1982). If not stated otherwise, strains were grown in liquid LB medium (Maniatis et al., 1982) or a modified M9 salt medium (Wehrmann et al., 2017) supplemented with 5 mM 2-phenylethanol, 5 mM 2-phenylacetaldehyde, 5 mM phenylacetic acid, or 25 mM succinate as carbon and energy source at 28°C to 30°C and shaking. 40 μg mL^-1^ kanamycin or 15 μg mL^-1^ gentamycin for *E. coli* and 40 μg mL^-1^ kanamycin, 20 μg mL^-1^ 5-fluorouracil, or 30 μg mL^-1^ gentamycin for *P. putida* strains was added to the medium for maintenance and selection, if indicated.

**Table 1:** Strains and plasmids used in the study

| Strains | Relevant features | Source or reference |
| --- | --- | --- |
| KT2440* | KT2440 with a markerless deletion of *upp*. Parent strain for deletion mutants. | (Graf and Altenbuchner, 2011) |
| $\Delta pedE$ | KT2440* with a markerless deletion of *pedE* | (Mückschel et al., 2012) |
| $\Delta pedA1A2BC$ | KT2440* with a markerless deletion of *pedA1A2BC* (PP_5538, PP_2669, PP_2668, PP_2667) | This study |
| $\Delta pedE \Delta pedA1A2BC$ | $\Delta pedE$ with markerless deletion of gene cluster *pedA1A2BC* | This study |
| $\Delta pedE \Delta pvdD$ | $\Delta pedE$ with a markerless deletion of *pvdD* (PP_4219) | This study |
| $\Delta pedH$ | KT2440* with a markerless deletion of *pedH* | (Mückschel et al., 2012) |
| $\Delta pedH \Delta pedA1A2BC$ | $\Delta pedH$ with a markerless deletion of gene cluster *pedA1A2BC* | this study |
| $\Delta pedE \Delta pedH$ | KT2440* with a markerless deletion of *pedE* and *pedH* | (Mückschel et al., 2012) |
| $\Delta pedE \Delta tatC1$ | $\Delta pedE$ with a markerless deletion of *tatC1* (PP_1039) | This study |
| $\Delta pedE \Delta tatC2$ | $\Delta pedE$ with a markerless deletion of *tatC2* (PP_5018) | This study |
| *E. coli* BL21 (DE3) | *F<sup>-</sup> ompT gal dcm lon hsdS<sub>B</sub>(r<sub>B</sub><sup>-</sup> m<sub>B</sub><sup>-</sup>) $\lambda$(DE3 [*lacI lacUV5-T7 gene 1 ind1 sam7 nin5*])* | (Studier and Moffatt, 1986) |
| *E. coli* TOP10 | *F<sup>-</sup> mcrA $\Delta$(mrr-hsdRMS-mcrBC) ϕ80lacZΔM15 ΔlacX74 nupG recA1 araD139 $\Delta$(ara-leu)7697 galE15 galK16 rpsL(Str<sup>R</sup>) endA1 λ<sup>-</sup>* | Invitrogen |
| *E. coli* HB101 | *F<sup>-</sup> mcrB mrr hsdS20(r<sub>B</sub><sup>-</sup> m<sub>B</sub><sup>-</sup>) recA13 leuB6 ara-14 proA2 lacY1 galK2 xyl-5 mtl-1 rpsL20(Sm<sup>R</sup>) gln V44 λ<sup>-</sup>* | (Boyer and Roulland-Dussoix, 1969) |
| *E. coli* PIR2 | *F Δlac169 rpoS(Am) robA1 creC510 hsdR514 endA reacA1 uidA(ΔMlui)::pir* | Invitrogen |
| *E. coli* CC118λpir | *$\Delta$(ara-leu) araD ΔlacX74 galE galK phoA20 thi-1 rpsE rpoB argE(Am) recA1 λpir phage lysogen* | (Herrero et al., 1990) |
| KT2440*::*Tn7M-pedH-lux* | KT2440* with insertion of *Tn7-M-pedH-lux* | This study |
| $\Delta pedA1A2BC$::*Tn7M-pedH-lux* | $\Delta pedA1A2BC$ with insertion of *Tn7-M-pedH-lux* | This study |
| KT2440*::*Tn7M-pedE-lux* | KT2440* with insertion of *Tn7-M-pedE-lux* | This study |
| **Plasmids** |  |  |
| pJOE6261.2 | Suicide vector for gene deletions | (Graf and Altenbuchner, 2011) |
| pMW10 | pJeM1 based vector for rhamnose inducible expression of *PedH* with C-terminal 6x His-tag | (Wehrmann et al., 2017) |
| pMW50 | pJOE6261.2 based deletion vector for gene *pvdD* (PP_4219) | This study |
| pMW57 | pJOE6261.2 based deletion vector for gene cluster *pedA1A2BC* | This study |
| pTn7-M | Km<sup>R</sup> Gm<sup>R</sup>, *ori R6K*, *Tn7L* and *Tn7R* extremities, standard multiple cloning site, *oriT* RP4 | (Zobel et al., 2015) |
| pRK600 | Cm<sup>R</sup>, *ori ColEI*, Tra<sup>+</sup> Mob<sup>+</sup> of RK2 | (Keen et al., 1988) |
| pTNS1 | Ap<sup>R</sup>, *ori R6K*, *TnSABC+D* operon | (Choi et al., 2005) |
| pSEVA226 | Km<sup>R</sup>, *ori RK2*, reporter vector harboring the *luxCDABE* operon | (Silva-Rocha et al., 2013) |
| pSEVA226-pedH | pSEVA226 with a *pedH-luxCDABE* fusion | This study |
| pSEVA226-pedE | pSEVA226 with a *pedE-luxCDABE* fusion | This study |
| pTn7-M-pedH-lux | pTn7-M with a *pedH-luxCDABE* fusion | This study |
| pTn7-M-pedE-lux | pTn7-M with a *pedE-luxCDABE* fusion | This study |

### Liquid medium growth experiments

Liquid growth experiments were performed in biological triplicates by monitoring the optical density at 600 nm (OD_600_) during growth in modified M9 medium supplemented with the corresponding carbon and energy sources (see above). For all experiments, washed cells from overnight cultures grown with succinate at 30°C and 180 rpm shaking were utilized to inoculate fresh medium with an OD_600_ of 0.01 to 0.05. Depending on the culture vessel, the incubation was carried out in 1 ml medium per well for 96-well 2 ml deep-well plates (Carl Roth) at 350 rpm shaking and 30°C, or 200 μL medium per well for 96-well microtiter plates (Sarstedt) at 180 rpm shaking and 28°C. If needed, different concentrations of LaCl_3_ (Sigma-Aldrich) were added to the medium.

### Construction of plasmids

The 600 bp regions upstream and downstream of gene *pvdD*, gene cluster *pedA1A2BC* and genes *tatC1* and *tatC2* were amplified from genomic DNA of *P. putida* KT2440 using primers pairs MWH56/MWH57 and MWH58/MWH59, MWH94/MWH95 and MWH96/MWH97, PBtatC1.1/PBtatC1.2 and PBtatC1.3/PBtaC1.4, and PBtatC2.1/PBtatC2.2 and PBtatC2.3/PBtatC2.4 to construct the deletion plasmids pMW50, pMW57, pJOE-tatC1 and pJOE-tatC2 (**Table 2**). The BamHI digested pJOE6261.2 as well as the two up- and downstream fragments were therefore joined together using one-step isothermal assembly (Gibson, 2011) and subsequently transformed into *E. coli* BL21(DE3) or TOP10 cells. Sanger sequencing confirmed the correctness of the plasmids.

**Table 2:**
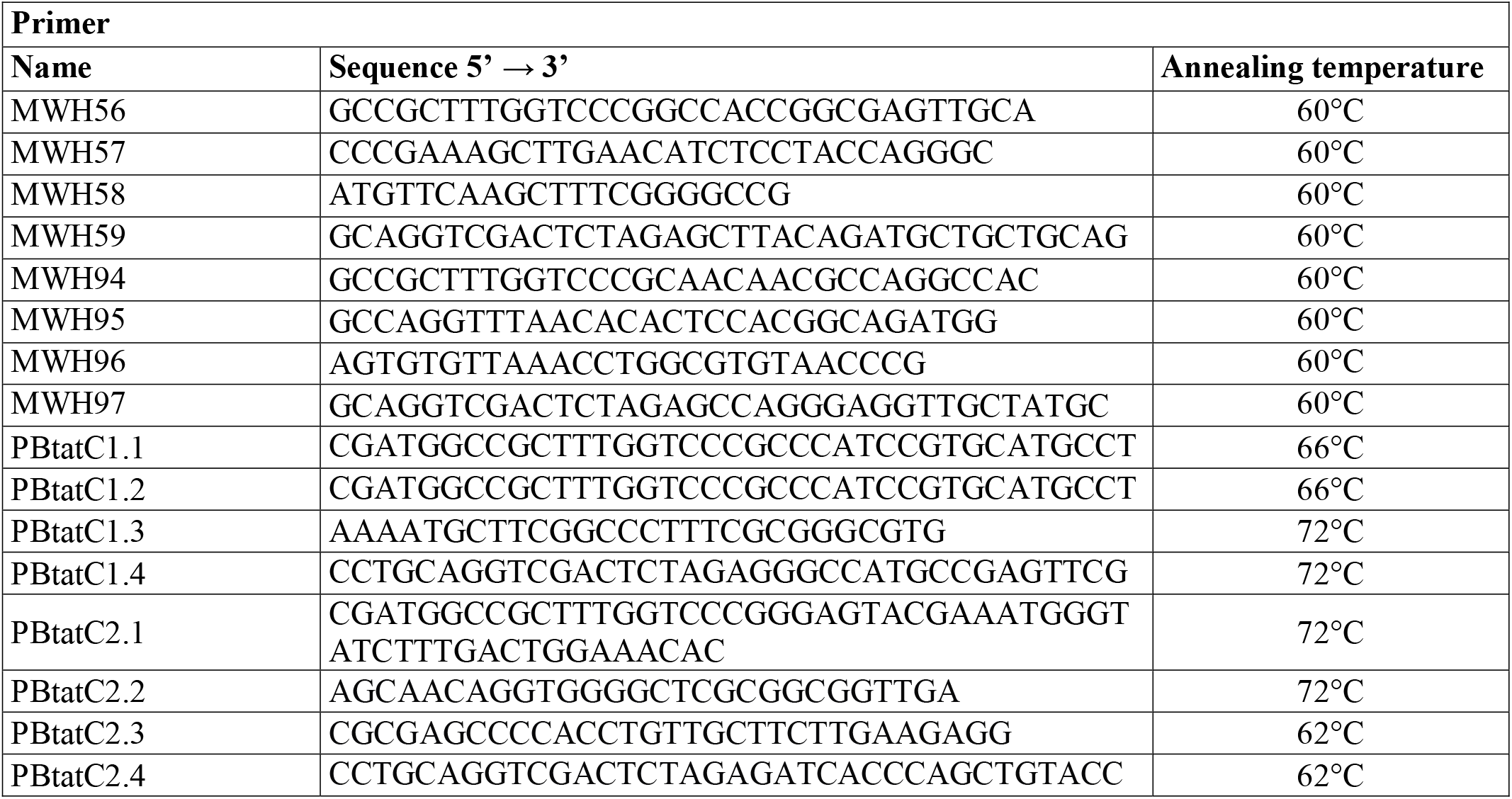
Primers used in the study

For measuring promoter activity of *pedE* and *pedH* in vivo, plasmids pTn7-M-pedH-lux and pTn7-M-pedE-lux were constructed. The DNA regions encompassing the promoters from *pedE* and *pedH* genes were amplified by PCR using the primer pairs p2674-FSac/p2674-RPst and p2679-FSac/p2679-RPst (Wehrmann et al., 2017). The PCR products were digested with SacI and PstI and inserted upstream the *luxCDABE* operon hosted by plasmid pSEVA226. The cargo module bearing the *pedE-lux* or *pedH-lux* fusion was then passed from the resulting pSEVA226-based constructs to pTn7-M as PacI/SpeI fragments.

### Strain constructions

For the deletion of chromosomal genes a previously described method for markerless gene deletions in *P. putida* KT2440 was used (Graf and Altenbuchner, 2011). In short, after transformation of the integration vectors carrying the up- and downstream region of the target gene, clones that were kanamycin (Kan) resistant and 5-fluorouracil (5-FU) sensitive were selected and one clone was incubated in liquid LB medium for 24 h at 30°C and 180 rpm shaking. Upon selection for 5-FU resistance and Kan sensitivity on minimal medium agarose plates, clones that carried the desired gene deletion were identified by colony PCR.

Integration of the pTn7-M based *pedH-lux* and *pedE-lux* fusions into the chromosome of *P. putida* KT2440 was performed by tetraparental mating using PIR2/pTn7-M-pedH-lux or PIR2/pTn7-M-pedE-lux as the donor, *E. coli* CC118 λpir/pTNS1 and *E. coli* HB101/pRK600 as helper strains and appropriate KT2440 strain as the recipient (Zobel et al., 2015). Briefly, cultures of the four strains grown under selective conditions were mixed, spotted on LB agar and incubated overnight at 28°C. Transconjugants were selected on cetrimide agar (Sigma-Aldrich) containing gentamicin. Correct chromosomal integration of mini-Tn*7* was checked by colony PCR using Pput-*glmS*DN and PTn*7*R primers as described elsewhere (Choi et al., 2005).

### Promoter activity assays

*P. putida* harboring a Tn7-based *pedH-lux* or *pedE-lux* fusion were grown overnight in M9 medium with 25 mM succinate, washed three times in M9 medium with no added carbon source, and suspended to an OD_600_ of 0.1 in the same medium with 1 mM 2-phenylethanol. For luminescence measurements, 198 μl of cell suspension was added to 2 μl of a 100-fold-concentrated metal salt solution in white 96-well plates with a clear bottom (μClear; Greiner Bio-One). Microtiter plates were incubated in a FLX-Xenius plate reader (SAFAS, Monaco) at 30°C with orbital shaking (600 rpm, amplitude 3 mm) and light emission and OD_600_ were recorded after the indicated time periods. Promoter activity was expressed as relative light units (RLU) normalized to the corresponding OD_600_. Experiments were performed in triplicates, and data are presented as the mean value with error bars representing the standard deviation.

## Results

*Pseudomonas putida* KT2440 makes use of a periplasmic oxidation system to grow on a variety of alcoholic substrates. Crucial to this system are two PQQ-dependent ethanol dehydrogenases (PQQ-EDHs), which share a similar substrate scope but differ in their metal cofactor dependency (Wehrmann *et al*., 2017). PedE makes use of a Ca^2+^-ion whereas PedH relies on the bioavailability of different rare earth elements (REE). During our studies, we found that the critical REE concentration that supports growth based on PedH activity differs dramatically depending on the minimal medium used. In a modified M9 medium, concentrations of about 10 μM of La^3+^ were necessary to observe PedH-dependent growth with 2-phenylethanol while only about 20-100 nM La^3+^ were required in MP medium (Wehrmann et al., 2017). One major difference between the two minimal media lies in their trace element composition and the respective metal ion concentrations (**Table 3**). The concentrations of copper, iron, manganese, and zinc are between 2x and 7x higher in the modified M9 medium compared to MP medium, and other trace elements such as boron, cobalt, nickel, or tungsten are only present in one out of the two media. To study the impact of the trace element solution (TES) on growth in the presence of La^3+^, we used the Δ*pedE* strain growing on 2-phenylethanol in M9 minimal medium in the presence and absence of TES.

**Table 3:**
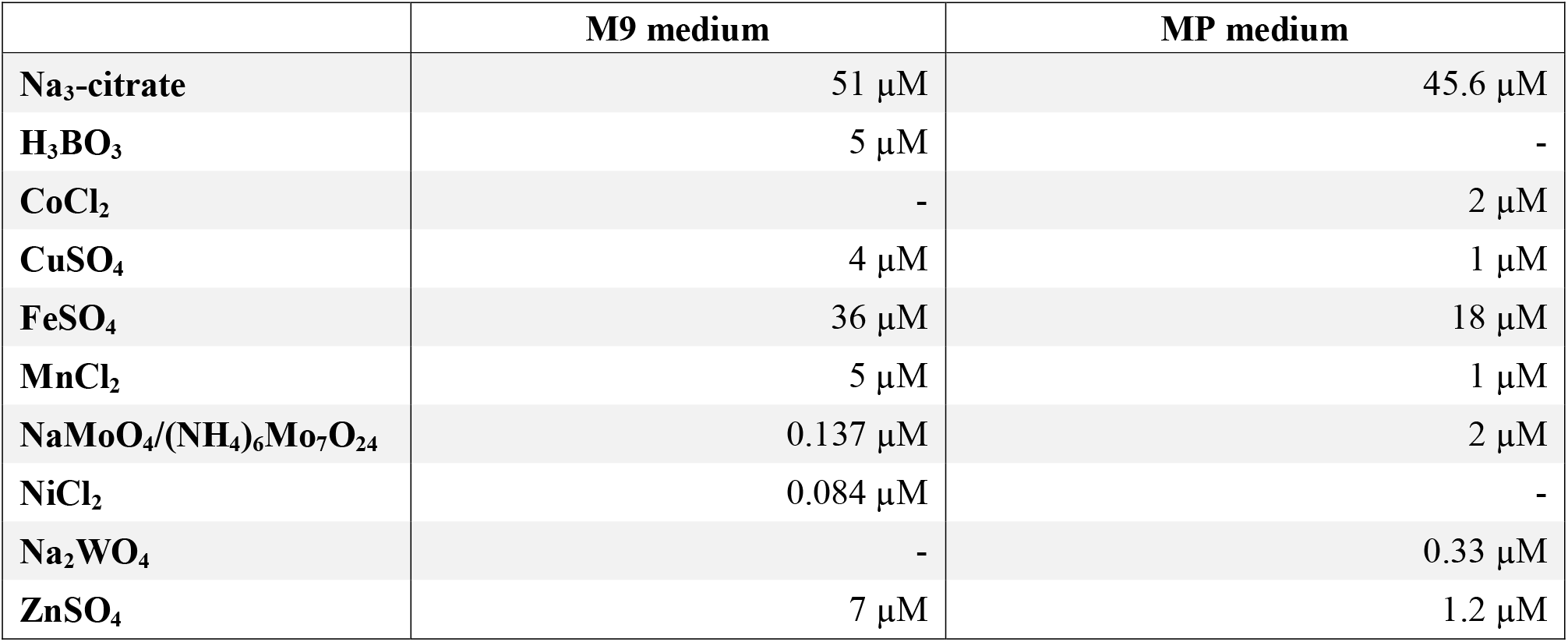
Trace element concentrations of M9 medium and MP medium.

While a critical La^3+^ concentration of 10 μM or higher was needed in the presence of TES to support PedH-dependent growth, this concentration dropped to as little as 10 nM La^3+^ in the absence of TES (**Figure 1A**). Similarly, inhibition of PedE-dependent growth by La^3+^ in strain Δ*pedH* differed dramatically depending on the presence of TES (**Figure 1B**). In the presence of TES, the addition of ≥ 100 μM of La^3+^ was required for growth inhibition in the Δ*pedH* strain within 48 h of incubation, whereas a minimum of only ≥ 1 μM La^3+^ was required in the absence of TES. From these experiments, we conclude that also a non-complemented minimal medium contains low, but sufficient, amounts of essential trace elements to allow growth even in the absence of TES. To find out whether the trace element mixture or a single trace element was causing the observed differences, we analyzed the growth of strain Δ*pedE* in more detail (**Figure 2A**). For concentrations of H_3_BO_3_, NaMoO_4_, NiSO_4_, and MnCl_2_ similar to those found in complemented M9 medium, PedH-dependent growth with 2-phenylethanol in the presence of 10 nM La^3+^ was observed. In contrast, upon the individual supplementation with 4 μM CuSO_4_, 36 μM FeSO_4_, or 7 μM ZnSO_4_ PedH-dependent growth could not be observed with 10 nM La^3+^, as it was the case upon the supplementation with TES. Since citrate is used as a metal chelator in TES, we further tested the impact of citrate on growth inhibition of CuSO_4_, FeSO_4_, and ZnSO_4_ when used as additional supplement (**Figure 2B**). The addition of 50 μM of Na_3_-citrate restored growth of the Δ*pedE* strain in the presence of 4 μM CuSO_4_ and 7 μM ZnSO_4_, even though cell growth was still impaired for Zn-containing medium. However, in cultures containing 36 μM FeSO_4_ the addition of citrate had no effect, strongly indicating that FeSO_4_ is predominantly responsible for the inhibition of PedH-dependent growth of the Δ*pedE* strain under low La^3+^ concentrations in a TES complemented M9 medium.

**Figure 1:**
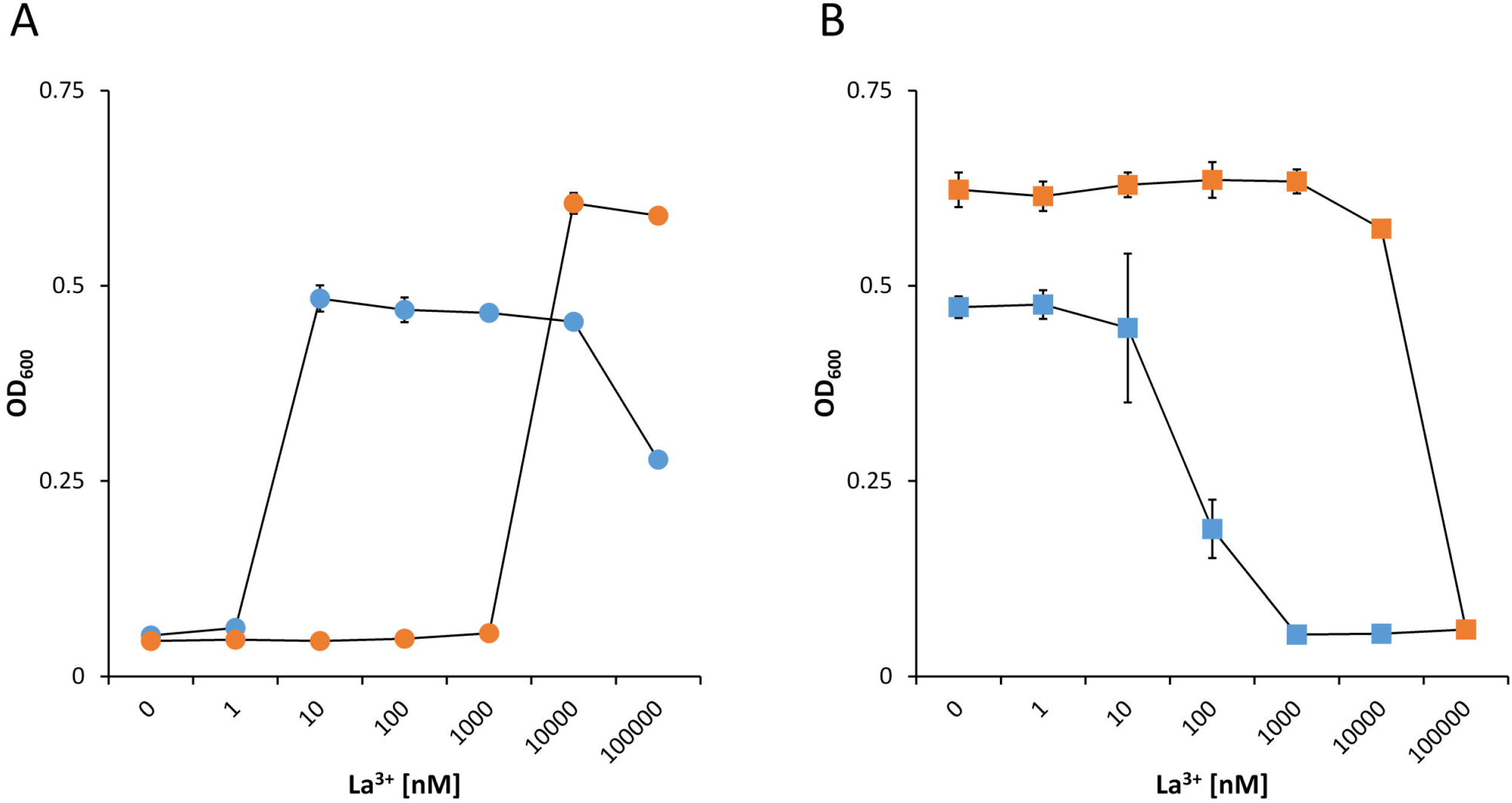
Growth of strain Δ*pedE* (**A**, dots) and Δ*pedH* (**B**, squares) in 1 mL liquid M9 medium in 96-well deep-well plates with 5 mM 2-phenylethanol and various concentrations of La^3+^ in the presence (orange) or absence (blue) of trace element solution (TES). OD_600_ was determined upon 48 h of incubation at 30°C and 350 rpm. Data are presented as the mean values of biological triplicates and error bars represent the corresponding standard deviations.

**Figure 2:**
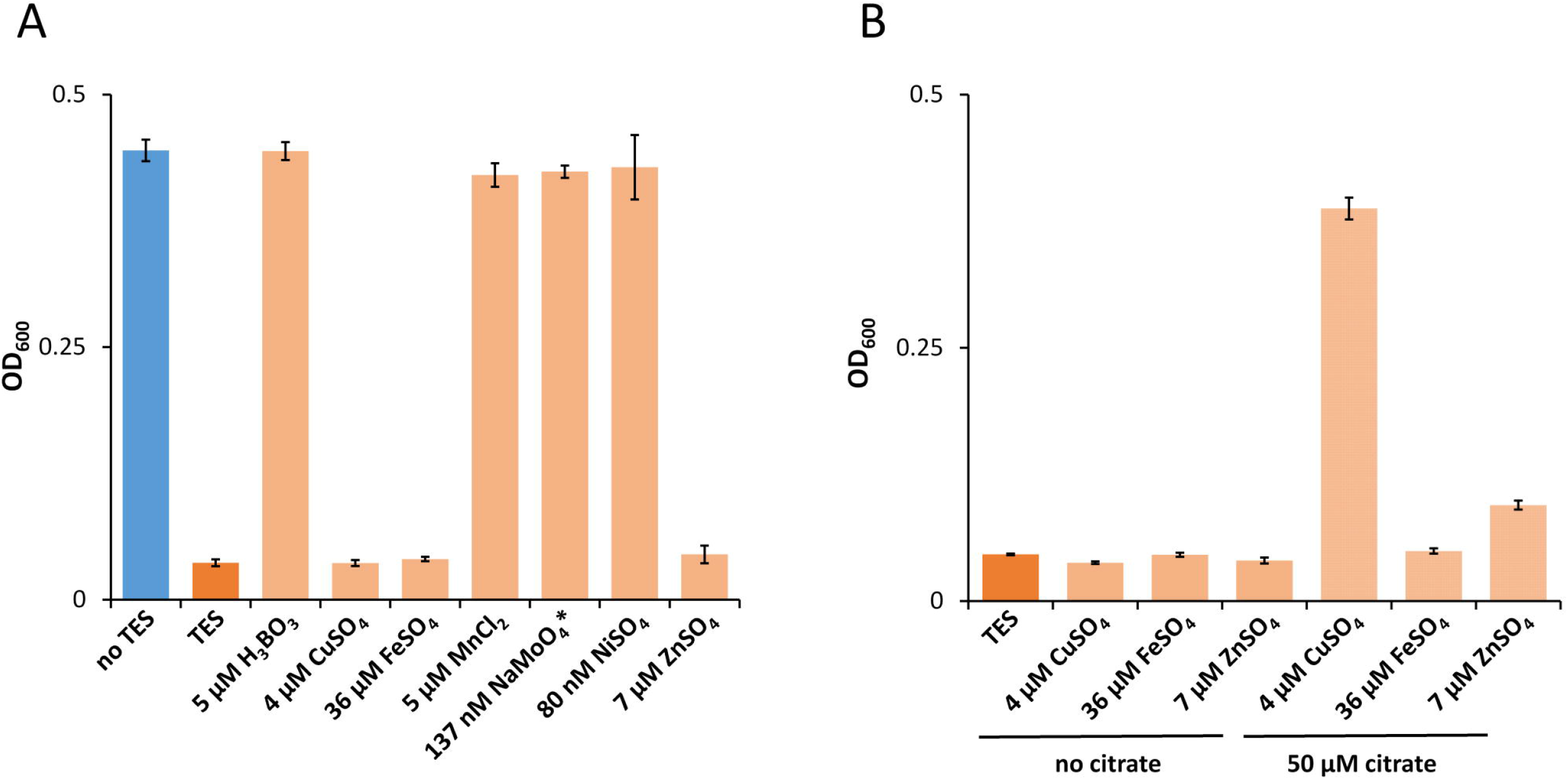
**A**) Growth of Δ*pedE* in 1 mL liquid M9 medium in 96-well deep-well plates with 5 mM 2-phenylethanol and 10 nM La^3+^ in the presence of trace element solution (TES) or individual components thereof. **B**) Growth of Δ*pedE* as described in **A** with and without the additional supplementation of 50 μM Na_3_-citrate. OD_600_ was determined upon 48 h of incubation at 30°C and 350 rpm. Data are presented as the mean values of biological triplicates and error bars represent the corresponding standard deviations. *Probes with 137 nM NaMoO_4_ also contain 5 μM H_3_BO_3_ and 84 nM NiSO_4_.

To acquire iron under restricted conditions, *P. putida* KT2440 can excrete two variants of the siderophore pyoverdine (Salah El Din et al., 1997). Beside their great specificity towards Fe^3+^, different pyoverdines can also chelate other ions including Al^3+^, Cu^2+^, Eu^3+^, or Tb^3+^, although with lower affinity (Braud et al., 2009a, 2009b). To test whether pyoverdine production in response to low iron conditions facilitates growth under low La^3+^ conditions, the mutant strain Δ*pedE* Δ*pvdD* was constructed. This strain is no longer able to produce the two pyoverdines due to the loss of the non-ribosomal peptide synthetase *pvdD* (PP_4219; formerly known as *ppsD*) (Matilla et al., 2007), which was confirmed upon growth on agar plates (**Figure 3C**). In experiments with varying FeSO_4_ supplementation, we found that PedH-dependent growth of strain Δ*pedE* was only observed for FeSO_4_ concentrations ≤ 10 μM under low (10 nM) La^3+^ conditions (**Figure 3A**). With ≥ 20 μM FeSO_4_ in the medium, no growth was observed. Strain Δ*pedE* Δ*pvdD* exhibited the same FeSO_4_-dependent growth phenotype as the parental strain under low La^3+^ concentrations. Under high (10 μM) La^3+^ conditions, strain Δ*pedE* exhibited PedH-dependent growth under any tested FeSO_4_ concentration (**Figure 3B**). Notably, strain Δ*pedE* Δ*pvdD* showed nearly the same growth pattern as Δ*pedE* under high La^3+^ concentrations, with the exception of the condition where no FeSO_4_ was added to the medium. Under this condition, no growth was observed.

**Figure 3:**
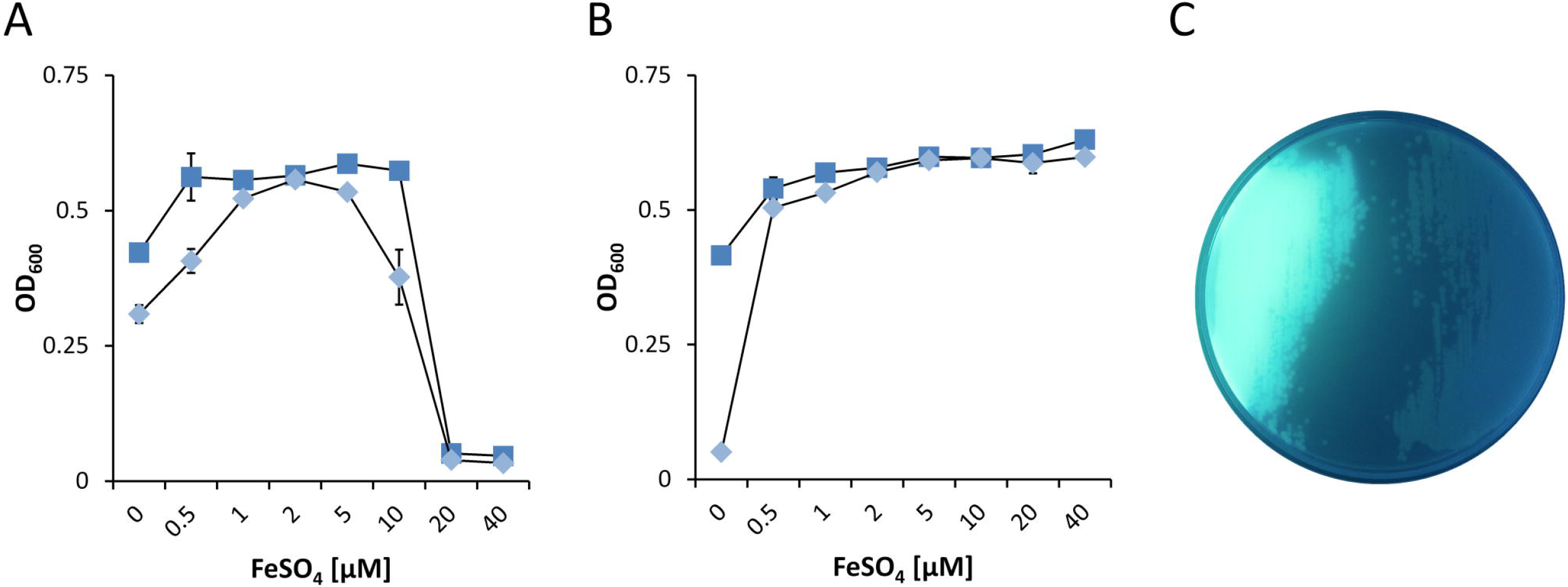
(**A** and **B**) Growth of Δ*pedE* (blue squares) and Δ*pedE* Δ*pvdD* (light blue diamonds) in 1 mL liquid M9 medium in 96-well deep-well plates without TES with 5 mM 2-phenylethanol and various concentrations of FeSO_4_ in the presence of 10 nM La^3+^ (**A**) or 10 μM La^3+^ (**B**). OD_600_ was determined upon 48 h of incubation at 30°C and 350 rpm. Data are presented as the mean values of biological triplicates and error bars represent the corresponding standard deviations. (**C**) Pyoverdine production by strains Δ*pedE* (left) and Δ*pedE* Δ*pvdD* (right) grown on cetrimide agar plates examined under blue light.

From these data, it can be speculated that beside the PedH-dependent growth also the inhibition of PedE-dependent growth is dependent on the FeSO_4_ to La^3+^ ratio. In the presence of 10 nM La^3+^, *pedE* promoter activity was comparably high and increased with increasing FeSO_4_ concentrations. In addition, strain Δ*pedH* grew readily on 2-phenylethanol under all these conditions even with no FeSO_4_ supplementation (**Figure 4A**). When 10 μM La^3+^ was available, no growth of the Δ*pedH* mutant was observed in presence of ≤ 20 μM FeSO_4_ and the *pedE* promoter activities were low (**Figure 4B**). However, when 40 μM FeSO_4_ were present in the medium, representing a 4-fold excess compared to La^3+^, PedE-dependent growth and an increased *pedE* promoter activity was detected.

**Figure 4:**
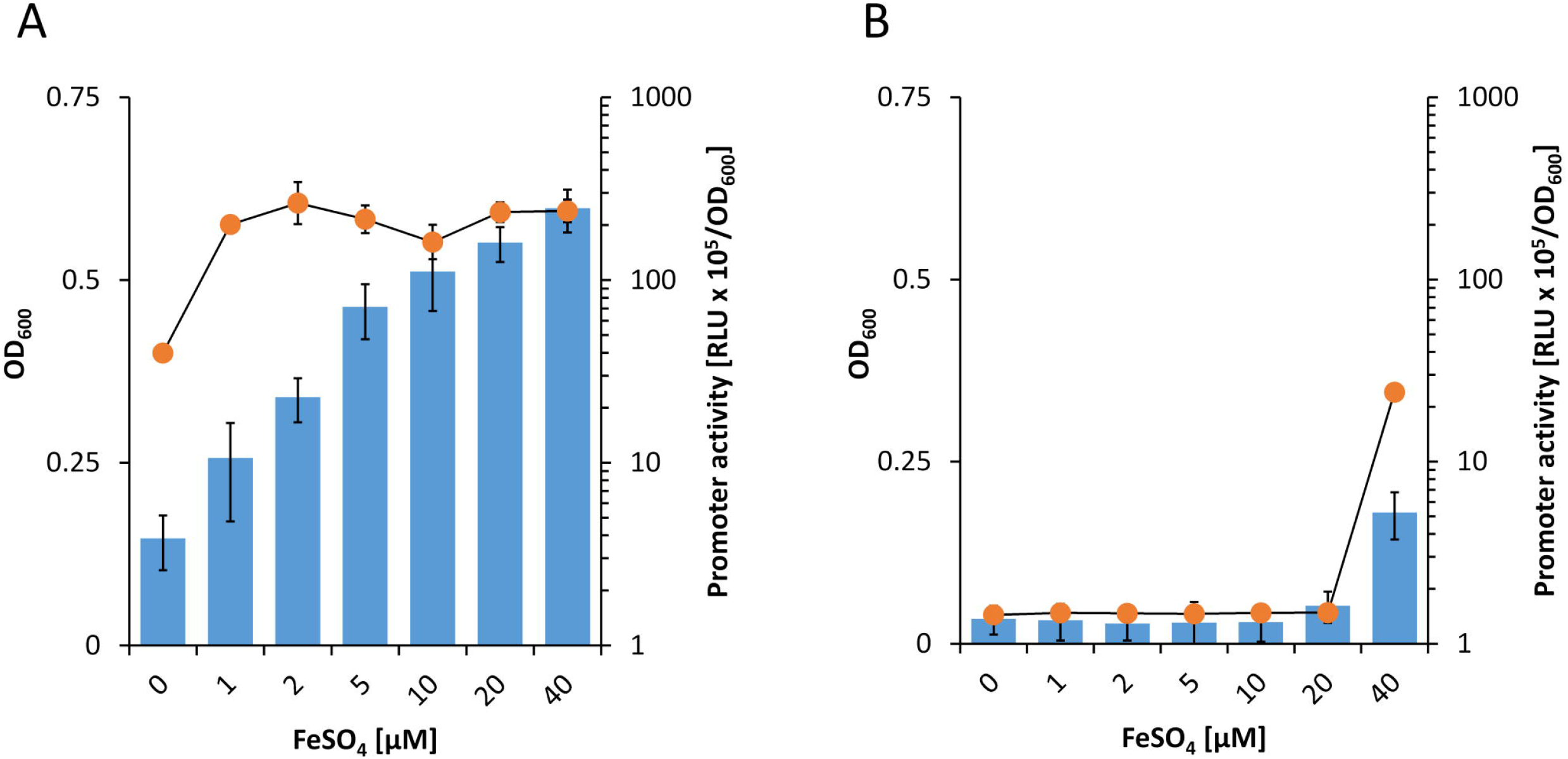
Activities of the *pedE* promoter (blue bars) in strain KT2440* during incubation in M9 medium with 2-phenylethanol, no TES and 10 nM (**A**) or 10 μM La^3+^ (**B**) as well as different FeSO_4_ concentrations. Promoter activities were determined upon 8 h of incubation at 600 rpm and 30°C. Growth of strain Δ*pedH* (orange dots) in M9 medium with 2-phenylethanol, no TES and 10 nM (**A**) or 10 μM (**B**) La^3+^ as well as different FeSO_4_ concentrations. Cells were incubated for 48 h in 96 deep-well plates at 30°C and 350 rpm prior to OD_600_ measurements. Data are presented as the mean values of biological triplicates and error bars represent the corresponding standard deviations.

Due to the very low concentrations of REEs (nM range) required for REE-dependent growth it is commonly speculated that specific REE uptake systems must exist. From our previous results, we can conclude that pyoverdine is not such a system. A search of the genomic context of the *ped* gene cluster identified a putative ABC transporter system located nearby the two PQQ-EDHs encoding genes *pedE* and *pedH* (**Figure 5A**). The ABC-transporter is predicted to be encoded as a single transcript by the online tool “Operon-mapper” (Taboada et al., 2018). It consists of four genes encoding a putative permease (*pedC* [PP_2667]), an ATP-binding protein (*pedB* [PP_2668]), a YVTN beta-propeller repeat protein of unknown function (*pedA2* [PP_2669]) and a periplasmic substrate-binding protein (*pedA1* [PP_5538]). While efflux systems are usually composed of the transmembrane domains and nucleotide binding domains, ABC-dependent import system additionally require a substrate binding protein for functional transport (Biemans-Oldehinkel et al., 2006). As the gene *pedA1* is predicted to be such a substrate binding protein, it is very likely that this transporter represents an import system.

**Figure 5:**
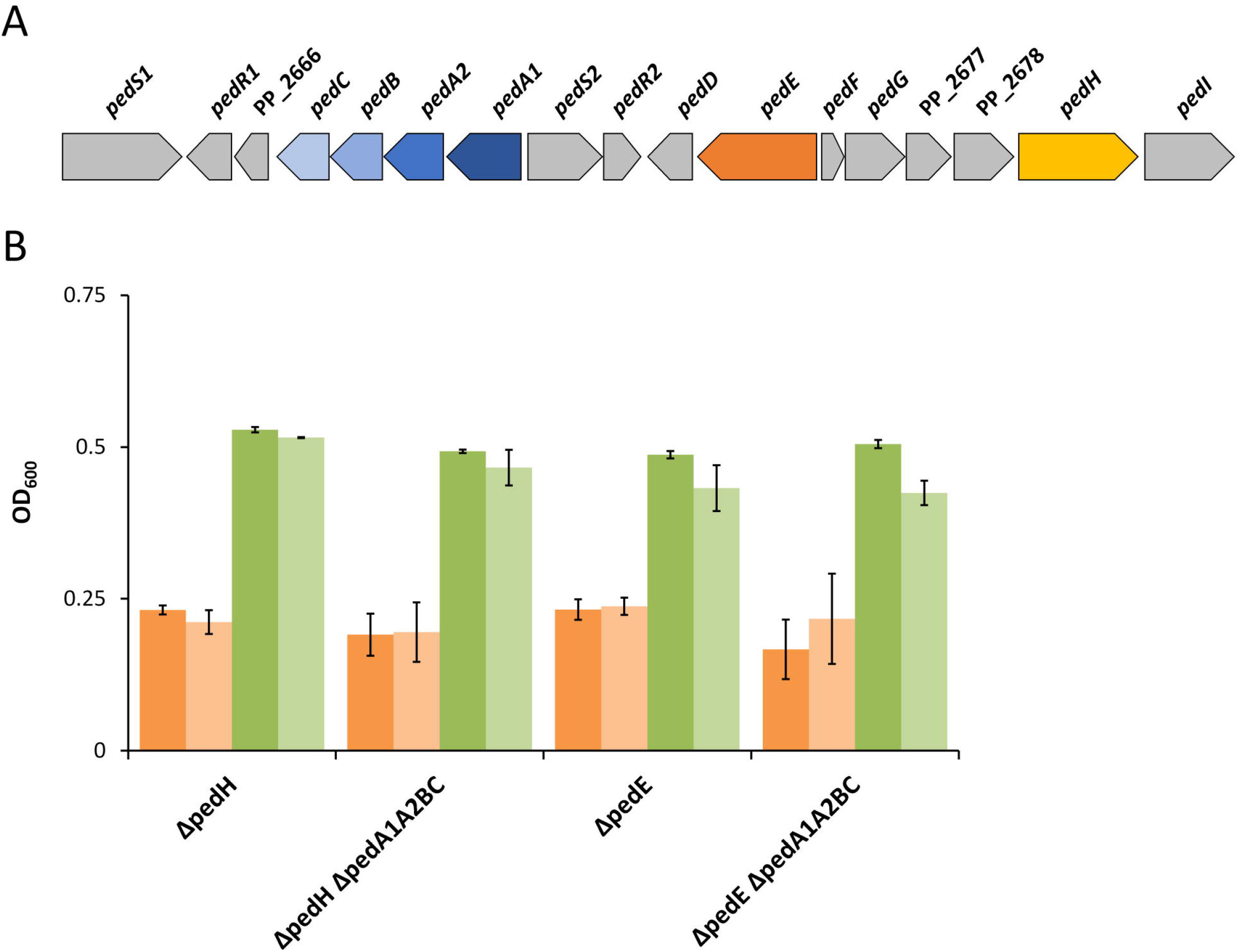
**A**) Genomic organization of the *ped* cluster in *P. putida* KT2440. Nomenclature in analogy to *P. putida* U as suggested by Arias *et al*.(Arias et al., 2008) **B**) Growth of Δ*pedH*, Δ*pedH* Δ*pedA1A2BC*, Δ*pedE*, and Δ*pedE* Δ*pedA1A2BC* strains in liquid M9 medium with TES and 5 mM phenylacetaldehyde (orange bars) or 5 mM phenylacetic acid (green bars) and either 0 μM La^3+^(dark green and dark orange bars) or 100 μM La^3+^ (light green and light orange bars). OD_600_ was determined upon 48 h of incubation at 30°C and 180 rpm. Data are presented as the mean values of biological triplicates and error bars represent the corresponding standard deviations.

ABC-dependent importers can be specific for carbon substrates or metal ions. Growth experiments with Δ*pedE*, Δ*pedH*, Δ*pedE* Δ*pedA1A2BC*, and Δ*pedH* Δ*pedA1A2BC* demonstrated that independent of La^3+^ (100 μM) availability, all strains were capable of growing with the oxidized degradation intermediates of 2-phenylethanol, namely 2-phenylacetaldehyde and phenylacetic acid, to a similar OD_600_ within 48 h of incubation (**Figure 5B**), indicating that the transport system is not involved in carbon substrate uptake.

When subsequently different La^3+^ concentrations were tested in a similar setup, we found that PedH-dependent growth on 2-phenylethanol of strain Δ*pedE* Δ*pedA1A2BC* was inhibited for the first 48 h of incubation under all concentrations tested, irrespectively of the presence or absence of TES (**Figure 6A**). This was in contrast to the Δ*pedE* deletion strain, which grew in the presence of ≥ 10 nM La^3+^ or ≥ 10 μM La^3+^ depending on TES availability (**Figure 1A**; **Figure 6A** pale symbols and lines). Upon an increased incubation time of 120 h, however, strain Δ*pedE* Δ*pedA1A2BC* eventually did grow with 1 and 10 μM La^3+^ or 100 μM La^3+^ depending on TES addition (**Figure 6B**). Notably, beside the substantial difference in lag-phase, also the critical REE concentration for PedH-dependent growth of Δ*pedE* Δ*pedA1A2BC* was increased by 100-fold compared to the Δ*pedE* strain under all conditions tested. Assuming that PedA1A2BC is specific for REE uptake, growth with the Ca^2+^-dependent enzyme PedE should not be influenced by a loss of the transporter function. When we tested the Δ*pedH* and Δ*pedH* Δ*pedA1A2BC* strain, we indeed could not find any difference in growth as both strains exhibited a similar inhibition pattern for concentrations ≥ 1 μM La^3+^ or ≥ 100 μM La^3+^ depending on the absence or presence of TES in the medium (**Figure 6C**).

**Figure 6:**
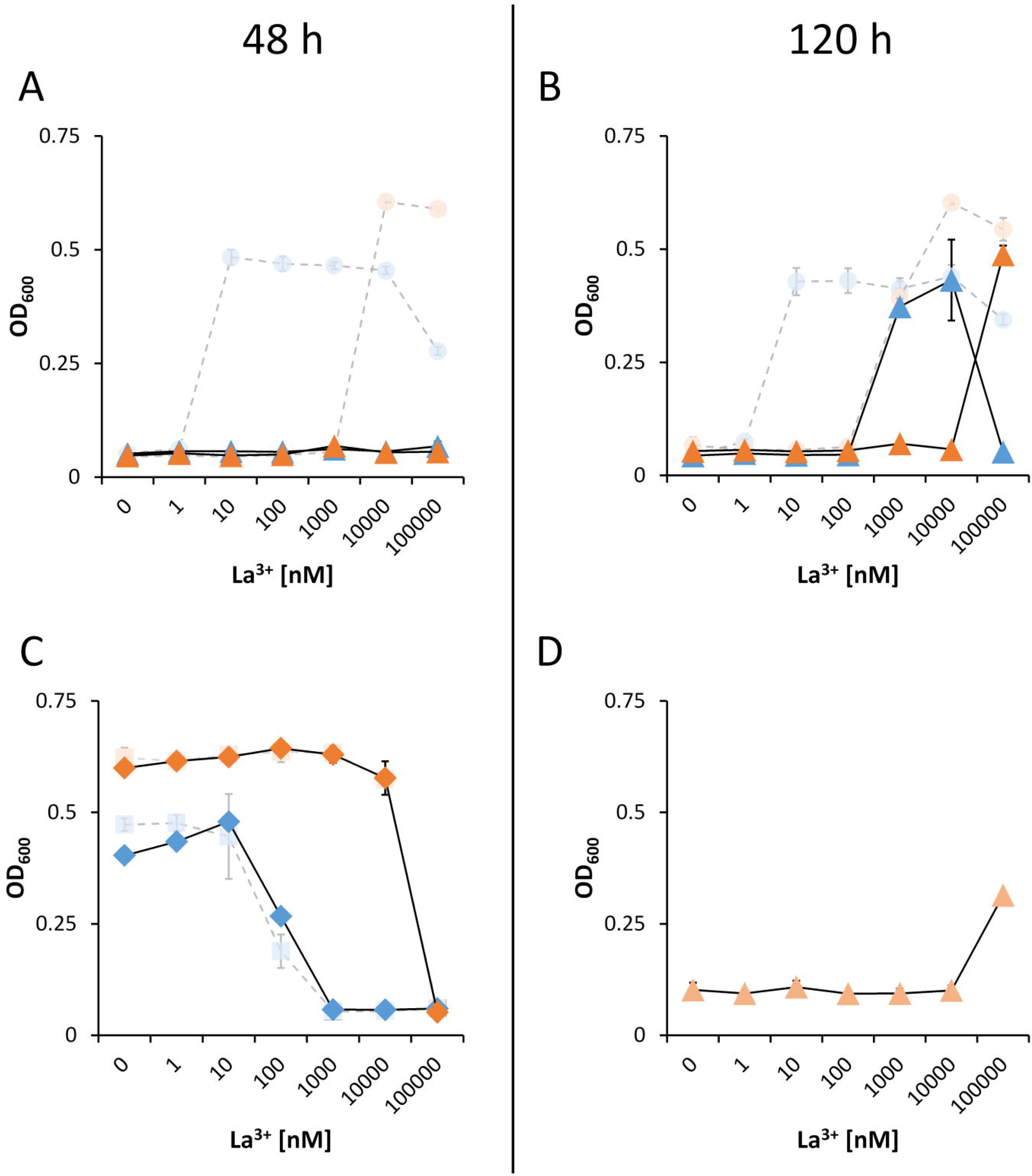
Growth of strains Δ*pedE* Δ*pedA1A2BC* (**A** and **B**, triangles) and Δ*pedH* Δ*pedA1A2BC* (**C**, diamonds) in liquid M9 medium with 5 mM 2-phenylethanol with (orange) or without (blue) TES and various concentrations of La^3+^. Pale circles and squares represent the growth of Δ*pedE* and Δ*pedH* parental strains. Growth of strains Δ*pedE* and Δ*pedH* in **A**) and **C**) represents the restated data from Figure 1 for better comparability. **D**) Growth of strain Δ*pedE* Δ*pedH* Δ*pedA1A2BC* harboring plasmid pMW10 (light orange triangles) in liquid M9 medium with 5 mM 2-phenylethanol, 20 μg/ml kanamycin, TES and various La^3+^ concentrations. OD_600_ was determined upon 48 h (**A** and **C**) or 120 h (**B** and **D**) of incubation at 30°C and 350 rpm. Data are presented as the mean values of biological triplicates, and error bars represent the corresponding standard deviations.

ABC-transporter systems, or the transported compounds, can be involved in transcriptional regulation of specific target genes (Biemans-Oldehinkel et al., 2006). Thus, the impaired growth under low La^3+^ concentrations of the Δ*pedE* Δ*pedA1A2BC* strain might be caused by the lack of transcriptional activation of the *pedH* gene. To test this hypothesis, strain Δ*pedE* Δ*pedH* Δ*pedA1A2BC* was complemented with a *pedH* gene independent of its natural promoter. Phenotypic analysis of this strain with 2-phenylethanol in the presence of TES and varying La^3+^ concentrations revealed no difference in the growth pattern when compared to strain Δ*pedE* Δ*pedA1A2BC* (**Figure 6D**). This indicated that the impaired growth phenotype of the ABC-transporter mutant is not due to a lack of transcriptional activation of *pedH*. To further validate this conclusion, *pedH* promoter activities were measured during incubation with 2-phenylethanol in strain Δ*pedA1A2BC* and its parental strain in the absence and presence of 10 μM La^3+^ (**Figure 7A**). Both strains showed a similar and more than 20 fold increased *pedH* promoter activity in response to La^3+^ supplementation (26-fold for KT2440*::Tn7-*pedH-lux* and 23-fold in Δ*pedA1A2BC*::Tn7-*pedH-lux*).

**Figure 7:**
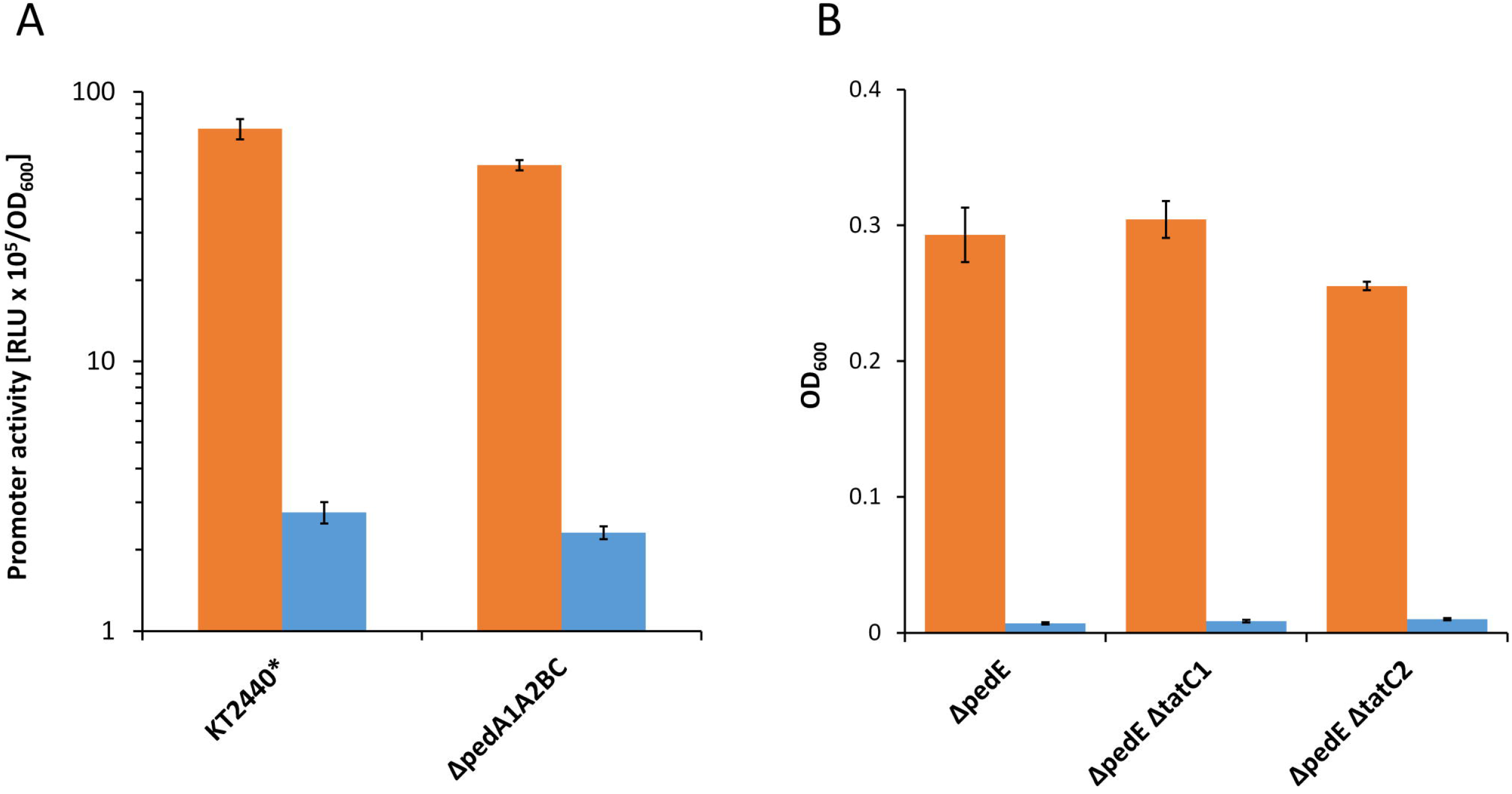
**A**) Promoter activities of the *pedH* promoter in the KT2440* and Δ*pedA1A2BC* background during incubation with 2-phenylethanol in presence (orange) or absence (blue) of 10 μM La^3+^. Promoter activities were determined upon 3 h of incubation at 600 rpm and 30°C. **B**) Growth of strain Δ*pedE* Δ*pedA1A2BC* on 2-phenylethanol in presence (orange) or absence (blue) of 10 μM La^3+^. OD_600_ was determined upon 48 h of incubation at 30°C and 180 rpm. Experiments were conducted in presence of TES. Data are presented as the mean values of biological triplicates and error bars represent the corresponding standard deviations.

In contrast to PedE, the signal peptide of PedH contains two adjacent arginine residues, which is an indication that it might be transported to the periplasm in a folded state via the Tat (twin-arginine translocation) protein translocation system (Berks, 2015). Therefore, one could argue that the transport of lanthanides into the cytoplasm might be beneficial as the incorporation into the active site of PedH could be more efficient during protein folding compared to the complementation of the apoenzyme in the periplasm. An initial analysis of the PedH signal peptide using different online software tools (TatP, PRED-TAT, SignalP 5.0, TatFind) could neither confirm nor refute this hypothesis (Almagro Armenteros et al., 2019; Bagos et al., 2010; Bendtsen et al., 2005; Rose et al., 2002). Therefore, we generated strains Δ*pedE* Δ*tatC1* and Δ*pedE* Δ*tatC2* in which the two individual TatC proteins (TatC1 [PP_1039] and TatC2 [PP_5018]) encoded in the genome of KT2440 are deleted. These strains should be restricted in the translocation of folded proteins into the periplasm, and if PedH would represent a Tat substrate, impaired growth on 2-phenylethanol in the presence of La^3+^ should be observable. However, neither *tatC1* nor *tatC2* mutation affected La^3+^-dependent growth on 2-phenylethanol (**Figure 7B**). Additionally, various attempts to generate the double *tatC1/C2* mutant strain were unsuccessful.

## Discussion

In the present study, we reveal that iron availability severely affects the REE-switch in *Pseudomonas putida* KT2440. This is evidenced by the reduction of the critical concentration of La^3+^ that is required both to promote PedH-dependent growth and for the repression of growth based on PedE activity. By using a Δ*pvdD* deletion strain, we demonstrate that the production of the iron chelating siderophore pyoverdine is not required for PedH-dependent growth under low La^3+^ conditions. Our data suggest that the observed effects during high Fe^2+/3+^/La^3+^ ratios are caused by mismetallation. In this scenario, the La^3+^-binding sites of proteins could be occupied by Fe^2+/3+^ ions that are in excess in the medium, and can also be present in the same 3+ oxidation state (Foster et al., 2014; Tottey et al., 2008; Tripathi and Srivastava, 2006; Webb, 1970). Transcriptional data show that *pedE* repression can be influenced by iron in a concentration dependent manner. Further, the impact of iron is not identical for PedE and PedH-dependent growth (100 fold vs. 1000 fold). Since PedE regulation is solely dependent on PedS2 (Wehrmann et al., 2017; 2018), these data are thus supportive of such a mismetallation hypothesis, assuming that the sensor histidine kinase PedS2 and PedH have different binding affinities to La^3+^ and/or Fe^2+/3+^.

The same hypothesis might similarly explain why under high La^3+^ concentrations in the absence of Fe^3+^ supplementation, a pyoverdine-deficient strain is strongly impaired in growth. In this scenario the Fe^2+/3+^ binding sites of pyoverdine-independent Fe transporters, such as the ferrichrome, ferrioxamine and ferric citrate uptake systems, might be occupied by La^3+^ and prevent binding of Fe^2+/3+^ ions (Cornelis, 2010; Jurkevitch et al., 1992). Consequently, a pyoverdine deficient strain would be unable to take up enough of this essential element that is, most likely, present at trace levels in the medium even without additional supplementation.

It is further interesting to point out that also micromolar Cu^2+^ and Zn^2+^ inhibited growth in presence of La^3+^ in the nanomolar range, although these metals do not exist in the same 3+ oxidation state under natural conditions. They are, however, the divalent transition metals that form the most stable complexes irrespective of the nature of the ligand, and as such also competitively bind non-cognate metal binding sites with high strength (Foster et al., 2014; Irving and Williams, 1953). Notably, Cu^2+^ has also been reported to interfere with REE-dependent regulation of PQQ-dependent methanol dehydrogenases in *M. trichosporium* OB3b (Gu et al., 2016; Gu and Semrau, 2017), and it is tempting to speculate that mismetallation might be involved in this process, too.

We provide compelling evidence that the predicted ABC-transporter PedA1A2BC is essential for PedH-dependent growth under low concentrations of La^3+^. Based on the PedE-dependent growth phenotype, we can further show that PedA1A2BC is not involved in transcriptional repression of *pedE* under low La^3+^-conditions. The fact that a Δ*pedE* Δ*pedA1A2BC* mutant strain can only grow with a 100-fold higher concentration of La^3+^ compared to the Δ*pedE* single mutant strongly indicates that PedA1A2BC functions as a La^3+^-specific importer into the cytoplasm. In very recent studies it was demonstrated that in several *Methylobacterium extorquens* strains a similar ABC-transporter system is required for Ln^3+^-dependent growth (Ochsner et al., 2019; Roszczenko-Jasinska et al., 2019). A BLAST analysis revealed that these ABC transporters show high similarities to all four genes of the *pedA1A2BC* operon (>43% sequence identity for *pedA2* and >50% for *pedA1, pedB* and *pedC*) and that all bacterial strains that have been reported to produce Ln^3+^-dependent PQQ-ADHs thus far, carry homologues of this transporter system in their genome. Using a protein-based fluorescent sensor with picomolar affinity for REEs, Mattocks *et al*.were able to demonstrate that *M. extorquens* indeed selectively takes up light REEs into its cytoplasm (Mattocks et al., 2019) and it was later shown that cytoplasmic REE-uptake depends on the presence of the previously identified ABC-transporter system (Roszczenko-Jasińska et al., 2019).

Since the PedH enzyme, like all currently known Ln^3+^-dependent enzymes, resides in the periplasm and since the purified apoenzyme of PedH can be converted into the catalytically active holoenzyme by Ln^3+^ supplementation *in vitro*, the question arises what the potential advantage of the postulated cytoplasmic Ln^3+^ uptake for *P. putida* would be. From our point of view, two different reasons can be imagined, namely that *i*) the REE-dependent PedH protein is folded within the cytoplasm and the incorporation of the La^3+^-cofactor is only possible or more efficient during the folding process; or *ii*) La^3+^ binds to a cytoplasmic protein that either represents a so-far uncharacterized transcriptional regulator or another REE-dependent enzyme. It has been demonstrated that the location of protein folding can regulate metal binding (Tottey et al., 2008). As such, Ln^3+^ insertion during folding in the cytoplasm, where metal concentrations are tightly regulated, could provide a means of preventing the Ln^3+^ binding site of PedH from mismetallation with potentially competitive binders such as Cu^2+^, Zn^2+^, or Fe^2+/3+^ in the periplasm. However, we could not find evidence that PedH is a Tat substrate and consequently transported into the periplasm as a folded protein (Berks, 2015), as the *tatC1* or *tatC2* mutants both still showed PedH-dependent growth. However, it cannot be excluded that the two Tat systems are functionally redundant since attempts to generate a *tatC1/C2* double mutant strain proved unsuccessful.

We can further conclude that the putative La^3+^ transport into the cytoplasm is not required to activate *pedH* transcription. It is however possible that additional genes/proteins required for PedH-dependent growth rely on, or are regulated by, the cytoplasmic presence of REEs. In this context it is interesting to note that in a recent proteomic approach, we found that besides PedE and PedH, additional proteins of unknown function show differential abundance in response to La^3+^ availability (Wehrmann et al., 2019). It will hence be interesting to find out in future studies whether any of these proteins is required for PedH function.

In *M. extorquens* PA1 and *M. extorquens* AM1, almost identical TonB-dependent receptor proteins (>99 % sequence identity) were found to be crucial for REE-dependent growth suggesting a specific Ln^3+^-binding chelator system in these organisms (Ochsner et al., 2019; Roszczenko-Jasinska et al., 2019). Interestingly, also in *Methylotuvimicrobium buryatense* 5GB 1C a TonB-dependent receptor was identified that is crucial for the REE-switch to occur (Groom et al., 2019). Interestingly, the latter receptor only shows < 20% sequence identity to those of the *M. extorquens* strains and could thus not have been identified by homology searches. In *P. putida* KT2440, no close homolog to any of the aforementioned TonB-dependent receptors could be identified (< 30% sequence identity). *P. putida* preferentially resides in the rhizosphere whereas *M. buryatense* and *M. extorquens* PA1 were isolated from the sediment of a soda lake (pH 9 – 9.5) and the phyllosphere of *Arabidopsis thaliana*, respectively (Kaluzhnaya et al., 2001; Knief et al., 2010). One could therefore speculate that a specific Ln^3+^-chelator system is perhaps not relevant in the rhizosphere due to the large reservoir of REEs within the soil and the usually acidic environment near the plant roots caused by the secretion of organic acids for phosphate solubilization (Raghothama and Karthikeyan, 2005; Ramos et al., 2016).The lack of a homologous TonB-dependent receptor could also be explained by structural differences in the REE-specific chelator system that might be employed by *P. putida* compared to that of the methylotrophic bacteria. Lastly, it is also possible that in *P. putida* the REE uptake across the outer membrane proceeds via the same chelator systems that are used for pyoverdin independent Fe-acquisition. This could further provide another explanation for the impact of the Fe^2+/3+^ to La^3+^ ratio on the REE-switch.

Overall, the present study expands the crucial role of a conserved ABC-transporter system, which was very recently identified as Ln^3+^-specific inner membrane transport system in methano- and methylotrophs, to non-methylotrophic organisms. It further provides new insight into the complexity of bacterial-metal interactions and demonstrates that Cu, Zn, and in particular Fe ions can strongly interfere with the REE-switch in *P. putida* most likely through mismetallation. The body of knowledge how REEs impact protein function, gene regulation, and consequently physiology of different microorganisms is rapidly increasing. As such, it will be very interesting to see when some of the most interesting questions, such as the cytoplasmic function of REEs or the nature and potential structural diversity of specific REE-chelator systems, will be resolved by future studies.

## Acknowledgements

The authors further declare no conflict of interest. Janosch Klebensberger and Matthias Wehrmann would like to thank Prof. Bernhard Hauer for his continuous support.

